# Three-dimensional imaging and single-cell transcriptomics of the human kidney implicate perturbation of lymphatics in alloimmunity

**DOI:** 10.1101/2022.10.28.514222

**Authors:** Daniyal J Jafree, Benjamin Stewart, Maria Kolatsi-Joannou, Benjamin Davis, Hannah Mitchell, Lauren G Russell, Lucía Marinas del Rey, William J Mason, Byung Il Lee, Lauren Heptinstall, Gideon Pomeranz, Dale Moulding, Laura Wilson, Tahmina Wickenden, Saif Malik, Natalie Holroyd, Claire Walsh, Jennifer C Chandler, Kevin X Cao, Paul JD Winyard, Karen L Price, Adrian S Woolf, Marc Aurel Busche, Simon Walker-Samuel, Peter J Scambler, Reza Motallebzadeh, Menna R Clatworthy, David A Long

## Abstract

Studies of the structural and molecular features of the lymphatic vasculature, which clears fluid, macromolecules and leukocytes from the tissue microenvironment, have largely relied on animal models, with limited information in human organs beyond traditional immunohistochemical assessment. Here, we use three-dimensional imaging and single-cell RNA-sequencing to study lymphatics in the human kidney. We found a hierarchical arrangement of lymphatic vessels within human kidneys, initiating along specialised nephron epithelium in the renal cortex and displaying a distinct, kidney-specific transcriptional profile. In chronic transplant rejection we found kidney allograft lymphatic expansion alongside a loss of structural hierarchy, with human leukocyte antigen-expressing lymphatic vessels infiltrating the medulla, presenting a putative target for alloreactive antibodies. This occurred concurrently with lymphatic vessels invading and interconnecting tertiary lymphoid structures at early stages of lymphocyte colonisation. Analysis of intercellular signalling revealed upregulation of co-inhibitory molecule-mediated CD4^+^ T cell-lymphatic crosstalk in rejecting kidneys, potentially acting to limit local alloimmune responses. Overall, we delineate novel structural and molecular features of human kidney lymphatics and reveal perturbations to their phenotype and transcriptome in the context of alloimmunity.

**SUMMARY:** Lymphatics regulate fluid balance and immune cell accumulation but are under-studied in human organs such as the kidney. Jafree and colleagues profiled human kidney lymphatics using three-dimensional imaging and single-cell RNA-sequencing, revealing structural and transcriptional perturbations in rejecting kidney transplants.

## INTRODUCTION

Lymphatics are blind-ended vessels that clear tissue fluid and macromolecules from the tissue microenvironment and, during inflammation, sprout to facilitate leukocyte clearance in a process termed lymphangiogenesis (Oliver et al. 2020; Petrova and Koh 2020; Stritt et al. 2021). Other lymphatic functions include absorption of dietary lipids (Bernier-Latmani and Petrova 2017), drainage of cerebrospinal fluid (Da Mesquita, Fu, et al. 2018), cholesterol transport (Ouimet et al. 2019), establishment of lung compliance (Jakus et al. 2014) and growth of organ progenitor niches (Peña-Jimenez et al. 2019; Gur-Cohen et al. 2019; Liu et al. 2020; Yoon et al. 2019). Accordingly, therapeutic manipulation of lymphatics has shown efficacy in animal models of lymphedema (Szőke et al. 2021; Yoon et al. 2003), myocardial infarction (Henri et al. 2016; Klotz et al. 2015; Vieira et al. 2018), malignancy (Hu et al. 2020; Song et al. 2020), neurodegeneration (Da Mesquita et al. 2021; Da Mesquita, Louveau, et al. 2018), and cystic kidney disease (Huang et al. 2016).

Despite these advances in animal studies, our understanding of lymphatic biology in human organs, and methods to target lymphatics clinically, are limited. Given the importance of tissue fluid composition in kidney physiology and function, a greater understanding of renal lymphatic in humans is needed. The human kidney contains epithelial nephrons which, allied with specialised capillary vasculature (Dumas et al. 2021; Jourde-Chiche et al. 2019), regulate plasma ultrafiltration, bodily fluid homeostasis, acid-base balance, blood pressure and several endocrine systems. A network of lymphatics exist within the kidney (Russell et al. 2019; Donnan et al. 2021; Jafree and Long 2020), present from at least the end of the first trimester in human fetal development and residing within the organ’s hilum and cortex (Jafree et al. 2019). From immunohistochemical studies, lymphangiogenesis has been recognised as a common feature of human kidney diseases (Heller et al. 2007; Sakamoto et al. 2009; Kerjaschki et al. 2004; Stuht et al. 2007; Tsuchimoto et al. 2017; Adair et al. 2007; Rodas et al. 2022). A more detailed assessment of human kidney lymphatic structure, molecular features and associations with other cell types may help to explain why, despite the postulated role of lymphatic expansion in the resolution of inflammation, immune infiltration, fibrosis and deterioration of organ function still occur in renal pathologies.

Here, we utilised immunostaining of large tissue samples to delineate the three-dimensional (3D) architecture of lymphatics in human kidneys, marrying this with single-cell RNA sequencing (scRNA-seq) to reveal the hierarchical arrangement of human kidney lymphatics and their organ-specific transcriptional features. We then interrogate chronic alloimmune rejection, featuring interstitial fibrosis and tubular atrophy; a leading cause of late allograft failure (Hariharan et al. 2021; Clayton et al. 2019) and associated with allograft lymphatic expansion with unknown consequences (Wong 2020). The alterations to lymphatic architecture we found in chronic rejection, occurring alongside defective trafficking of and molecular crosstalk with effector T cells, together implicate lymphatic dysfunction as a hallmark of alloimmunity with potential therapeutic implications.

## RESULTS AND DISCUSSION

### Characterisation of lymphatic vessel architecture, spatial relationships and molecular profile in the human kidney

We first characterised kidney lymphatic architecture in humans by obtaining kidney tissue from four deceased organ donors (**Table S1A**). Tissues were immunolabelled with a D2-40 monoclonal antibody targeting the transmembrane protein, podoplanin (PDPN) (Kahn et al. 2002), before confocal or lightsheet fluorescence microscopy (LSFM); techniques used to visualise kidney lymphatics during organogenesis (Jafree et al. 2019) and in murine disease models (Liu et al. 2021). PDPN^+^ vessel networks were discriminated in intact volumes of human kidney cortex up to 3 mm^3^ (**Fig.1A**). Mapping vessel radius revealed the structural hierarchy of human kidney lymphatics, with the smallest vessels at the initiation of the lymphatic network (radius ∼3.5 μm) arising in the cortex in confluence with larger vessels (radius ∼50 μm) at the corticomedullary junction (**Fig.1B**). These vessels co-stained for the canonical lymphatic marker (Banerji et al. 1999; Wigle and Oliver 1999) the transcription factor prospero homeobox protein 1 (PROX1), with nuclear PROX1^+^ localisation evident within PDPN^+^ lymphatics (**Fig.1C**). However, expression of the transmembrane glycoprotein lymphatic vessel endothelial hyaluronan receptor 1 (LYVE1) was sparse and limited to lymphatic vessel tips (**Fig.1D**).

**Figure 1.**
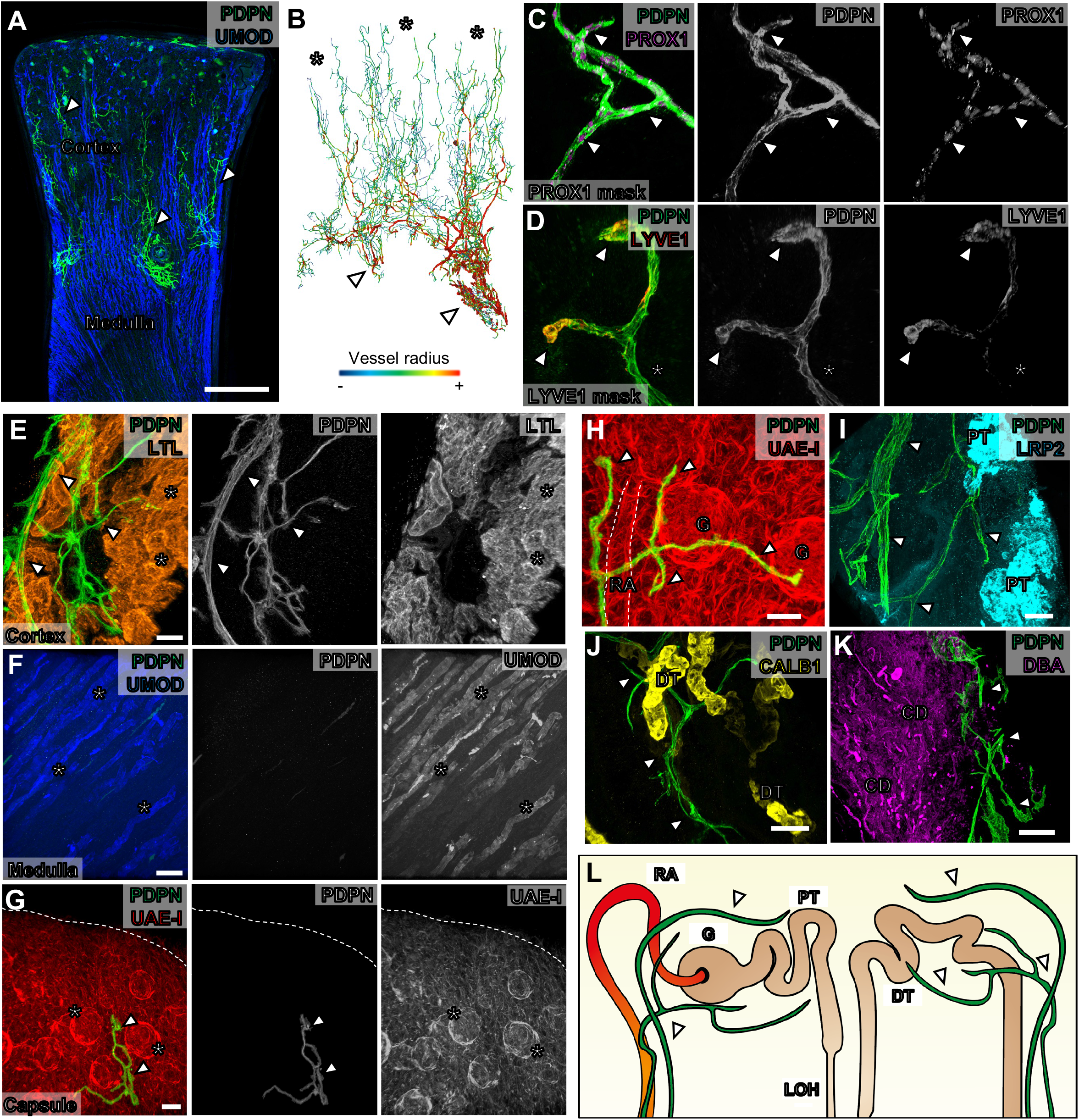
Three-dimensional imaging of lymphatics and their spatial relationships in the human kidney. **(A)** Representative maximum intensity *z*-projection, from low-resolution confocal tile scans, of *n =* 3 human kidney tissues labelled for podoplanin (PDPN) and uromodulin (UMOD), demonstrating PDPN^+^ lymphatics (arrowheads). Scale bar = 2000 μm. **(B)** Segmented and rendered LSFM imaging of lymphatics from the same kidney tissue in **A**, representative of *n =* 3 images. 3D colour renderings represent vessel branch radii, with blue representing the smallest radius (< 3.5 μm, asterisks) and red representing the largest radii (> 18 μm, arrowheads). **(C-D)** Representative 3D reconstruction of cortical regions from *n =* 2 human kidney tissues labeled for PDPN and either prospero homeobox protein 1 (PROX1) or lymphatic vessel endothelial hyaluronan receptor 1 (LYVE1). The PROX1 and LYVE1 signals are masked to only include expression from within the vessel, demonstrating expression of PDPN^+^ cells. Sparse membrane localization of LYVE1 is demonstrated (arrowheads). Representative of five regions of interest imaged. Scale bars = 30 μm. **(E-G)** Regional localization of lymphatics (arrowheads) in the human kidney using *Lotus tetragonolobus* lectin (LTL, cortex), UMOD (medulla) and *Ulex europaeus* agglutinin I (UEA-I, dotted lined delineating the capsule). Regional structures are indicated with asterisks, including proximal tubules in **E**, loops of Henle in **F** and glomeruli in **G**. Scale bars = 70 μm **(E)**, 150 μm **(F)**, 100 μm **(G). (H-K)** Spatial relationships of lymphatics (arrowheads) relative to UAE-I^+^ renal arterioles (RA) and glomeruli (G) in **H**, megalin (LRP2)^+^ proximal tubules (PT) in **I**, calbindin (CALB1)^+^ distal tubules (DT) in **J** and *Dolichos biflorus* agglutinin (DBA)^+^ collecting ducts (CD) in **K**. Scale bars = 50 μm **(H)**, 80 μm **(I** and **J)**, 300 μm **(K). (L)** Schematic depicting the spatial relationships of lymphatics (arrowheads) to nephron segments. All imaging from **E-K** representative of five regions of interest imaged across *n =* 2 kidneys.

We then assessed the precise anatomical location of lymphatic vessels relative to differentiated renal structures in the human kidney. Counterstaining to delineate nephron tubular epithelia, segments, including *Lotus tetragonolobus* lectin (LTL), which binds proximal tubular apical membranes within the cortex, and uromodulin (UMOD), expressed by loop of Henle epithelium in the medulla, confirmed PDPN^+^ blind-ended lymphatics in the renal cortex (**Fig.1E**) and their absence in the medulla (**Fig.1F**), consistent with previous studies (Ishikawa et al. 2006; Sakamoto et al. 2009; Kerjaschki et al. 2004; Stuht et al. 2007; Adair et al. 2007; Tsuchimoto et al. 2017). Lymphatics have been described in the sub-capsular space around the kidney (Holmes et al. 1977; Russell et al. 2019), but this was not evident in our samples, despite the capsule being left intact (**Fig.1G**). Within the renal cortex, PDPN^+^ lymphatics follow the course of UAE-I^+^ arterioles *en route* to glomeruli (**Fig.1H**); the site of plasma ultrafiltration, before giving off terminal branches adjacent to major sites of reabsorption of solute, ions and drugs, including megalin (LRP2)^+^ proximal tubules (**Fig.1I**) and calbindin 1 (CALB1)^+^ distal tubules (**Fig.1J**). Lymphatics then continue towards the kidney hilum adjacent to medullary *Dolichos biflorus* agglutinin (DBA)^+^ collecting ducts (**Fig.1K**). A model of these findings is presented in **Fig.1L**.

Lymphatics represent a rare cell type within the kidney. Therefore, to generate molecular profiles of human kidney lymphatics, we integrated published scRNA-seq data from 59 kidneys with additional data generated from five further samples (**Fig.2A**). This integrated kidney cell atlas contained 217,411 human kidney cells, including 151,038 ‘control’ cells from living donor biopsies or non-tumorous regions of tumour nephrectomies and 66,373 cells from diseased samples, including chronic kidney disease (CKD) and different aetiologies of transplant rejection (**Fig.S1A**). We identified 37 transcriptionally distinct clusters (**Fig.S1B**), including a lymphatic cluster (*n =* 700) distinct from five other blood endothelial clusters. We first examined cells from control samples to curate a lymphatic transcriptional signature, comprising 227 differentially expressed genes (DEG), that included those found within several relevant gene ontology (GO) gene-sets, including lymphatic fate commitment (GO:0060838, fold-enrichment > 100, *p =* 1.66 × 10^−2^) and lymphangiogenesis (GO:0001946, fold-enrichment = 67.4, *p =* 8.37 × 10^−3^). Canonical lymphatic markers including *PROX1* (log_2_FC = 2.97), *PDPN* (log_2_FC = 2.65), neuropilin 2 (*NRP2*, log_2_FC = 2.73) and chemokine (C-C motif) ligand (*CCL*)*21* (log_2_FC = 7.23) were among the top 20 DEGs in the LEC cluster (**Fig.2B**). Several postulated biomarkers of kidney disease (He et al. 2019; Tanaka et al. 2018; Shi et al. 2020) not previously known to be expressed by kidney lymphatics were identified, such as fatty acid binding protein 4 *(FABP4*, log_2_FC = 5.69) trefoil factor 3 (*TFF3*, log_2_FC = 5.58) and angiopoietin 2 (*ANGPT2*, log_2_FC = 2.46) (**Fig. 2B**).

**Figure 2.**
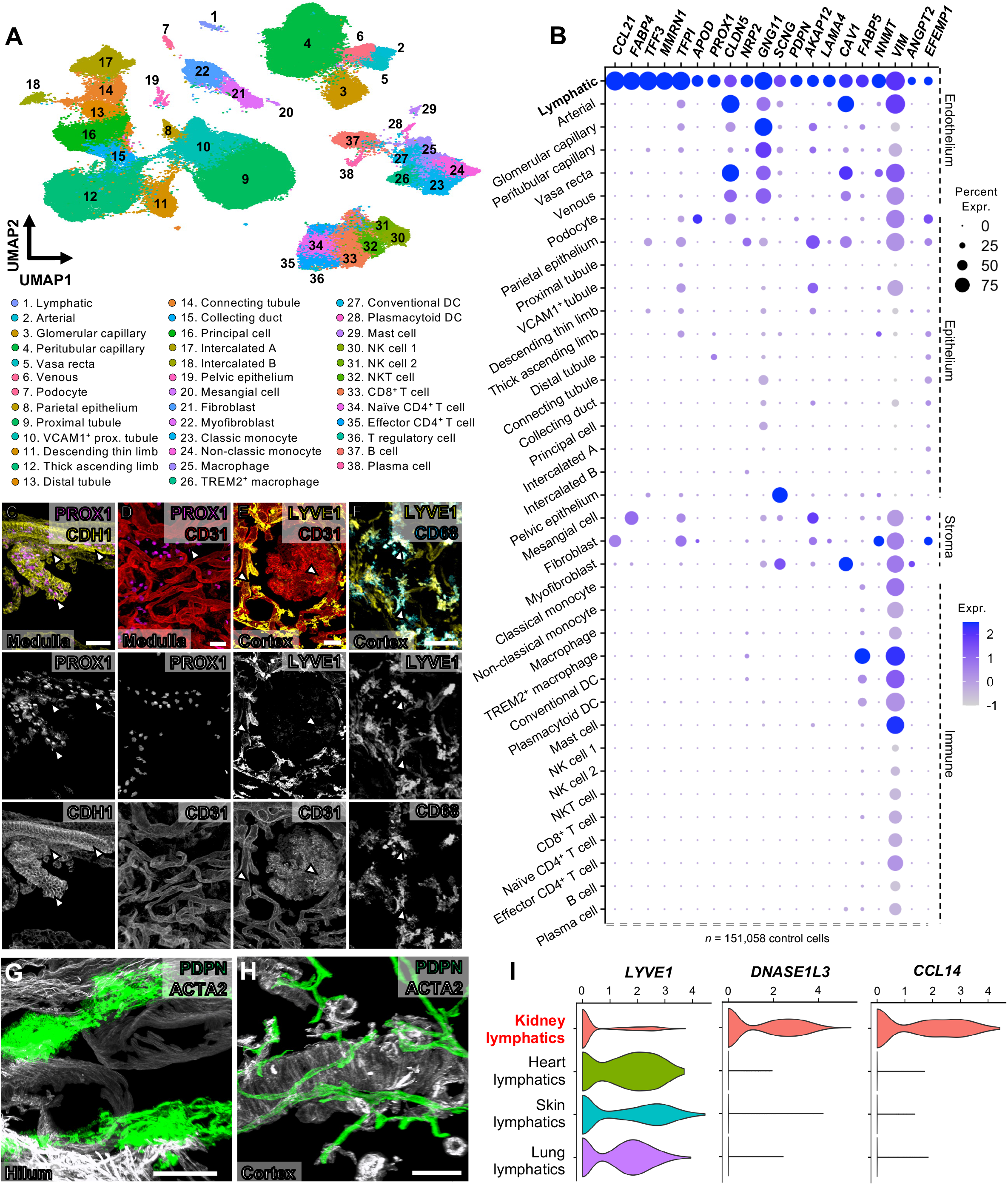
Profiling lymphatics and their molecular markers through single-cell RNA-sequencing of the human kidney. **(A)** Uniform manifold approximation and projection (UMAP) of an integrated atlas of 217,411 cells, including 151,058 ‘control’ cells from live biopsies or nephrectomies, 46,540 cells from different aetiologies of transplant rejection and 19,813 chronic kidney disease. DC, dendritic cell; NK, natural killer; TREM, triggering receptor expressed on myeloid cells 2; VCAM1, vascular cell adhesion protein 1. **(B)** Dot plot of top 20 markers of lymphatics profiled across all ‘control’ cell types in the atlas. Categorisation for each cell type is shown on the right. **(C-F)** Analysis of non-lymphatic expression of PROX1 and LYVE1 using 3D imaging. Arrowheads the show the expression of each marker relative to cadherin 1 (CDH1)^+^ medullary tubules **(C)**, CD31^+^ vasa recta **(D)** or peritubular capillaries **(E)** and CD68^+^ macrophages **(F)**. Scale bars = 50 μm (**C, E, F**), 30 μm (**D**). **(G-H)** Examination of α-smooth muscle actin (ACTA2) expression relative to PDPN^+^ lymphatics (arrowheads) in the renal hilum (**G**) and cortex (**H**). Scale bars = 50 μm (**G**), 100 μm (**H**). (**I**) Cross-organ comparison of lymphatics in the kidney, heart, skin and lung, represented by violin plots. Compared to other organs, kidney lymphatics express significantly lower *LYVE1* (log_2_FC = -1.21, *p* = 3.37 × 10^−20^) and higher deoxyribonuclease 1L3 (*DNASE1L3;* log_2_FC = 2.51, *p* = 2.33 × 10^−39^) and chemokine (C-C motif) ligand 14 (*CCL14;* log_2_FC = 3.12, *p* = 6.83 × 10^−92^) expression.

PROX1 and LYVE1 have been used to identify or target lymphatics in mouse studies of kidney disease (Donnan et al. 2021; Jafree and Long 2020) so we performed a detailed analysis of these markers in the human kidney. Our scRNA-seq data showed that *PROX1* expression was not limited to kidney lymphatics but was also evident in loop of Henle and distal convoluted tubule cells (**Fig.2B**). PROX1 expression was confirmed in E-cadherin (CDH1)^+^ medullary tubules (**Fig.2C**), mirroring observations in mouse (Kim et al. 2015), but was not detected in vasa recta endothelial cells at transcript (**Fig.2B**) or protein level (**Fig.2D**), in contrast to murine studies (Kenig-Kozlovsky et al. 2018; Liu et al. 2021). LYVE1 was expressed by renal macrophages (**Fig.2E**) and blood vasculature (**Fig.2F**), as noted previously in both mouse (H.-W. Lee et al. 2011) and human kidney (Marshall et al. 2022). These findings demonstrate PROX1 and LYVE1 expression beyond lymphatics in the human kidney; relevant for experiments using these markers for imaging or therapeutic manipulation.

Lymphatics possess phenotypic heterogeneity, with distinction between blind-ended capillaries lacking mural cell coverage and valve-containing smooth muscle-lined collecting vessels. Akin to blood vessels, different organs may also contain molecularly and structurally distinct lymphatics (Petrova and Koh 2018; Wong et al. 2018). To investigate these properties, we first stained kidney lymphatics for α-smooth muscle actin, and found, akin to lung (Reed et al. 2019) but unlike dermis or mesentery (Wang et al. 2017), that these vessels lack smooth muscle coverage (**Fig.2G-H**). To further assess inter-organ heterogeneity and determine if kidney lymphatics have a distinct molecular profile, we compared the transcriptome of human kidney lymphatics to lymphatics isolated from published scRNA-seq data of human heart, lung and skin (**Fig.2I**). Two of the top markers of kidney lymphatics, relative to lymphatics in other organs, included deoxyribonuclease (*DNASE*)*1L3* (log_2_FC = 2.51, *p* = 2.33 × 10^−39^) and the chemokine *CCL14* (log_2_FC = 3.12, *p* = 6.83 × 10^−92^). Notably, expression of *LYVE1* in kidney lymphatics was significantly lower than in lymphatics of other organs (log_2_FC = -1.21, *p* = 3.37 × 10^−20^) (**Fig.2I**). *LYVE1*, was only detectable in a quarter of kidney lymphatic cells, consistent with the spatial restriction of LYVE1 observed in 3D imaging (**Fig.1D**).

Taken together, our 3D imaging and scRNA-seq analyses support the organ-specificity of lymphatic structure and profile, extending these observations to the human kidney. We show human kidney lymphatics, which possess a capillary-like nature, initiate adjacent to specialised tubular epithelium in the cortex, and have a unique transcriptomic profile compared with other renal cells or lymphatics within other organs.

### Structural phenotype and origin of kidney lymphatics in chronic transplant rejection and their putative targeting by alloantibodies

Several seminal studies have demonstrated expansion of intra-graft lymphatics during allograft rejection in rodent models (Palin et al. 2013; Vass et al. 2012; Lin et al. 2021; Motallebzadeh et al. 2012) and in humans (Adair et al. 2007; Kerjaschki et al. 2004; Stuht et al. 2007; Tsuchimoto et al. 2017), suggesting that these vessels are either insufficient to resolve allograft inflammation or themselves partake in the process of rejection (Wong 2020). To probe this further, we used 3D lymphatic imaging and scRNA-seq to study kidneys with chronic rejection, which frequently involves both T cell and antibody-mediated alloimmune responses, leading to allograft injury (Loupy et al. 2022; Callemeyn et al. 2022). In three allograft explants with chronic mixed (T cell- and antibody-mediated) rejection (**Table S1B**) and using organ donor tissues as controls, we found disorganisation of the lymphatic network, with loss of structural hierarchy (**Fig.3A**), a seven-fold increase in lymphatic density (95.12 ± 49.21 *vs*. 690.3 ± 121.6 vessels / mm^3^, *p* = 0.0014) and infiltration of lymphatics into the kidney medulla (**Fig.3A-C**), which is normally devoid of these vessels (Donnan et al. 2021; Jafree and Long 2020; Russell et al. 2019). Additional abnormalities observed in rejection included a reduction in the mean vessel radius (10.84 ± 6.76 *vs*. 6.27 ± 3.95 μm, *p <* 0.0001) and branching angle (112.3 ± 28.87 *vs*. 101.4 ± 36.70°, *p* < 0.0001) compared to control kidneys.

**Figure 3.**
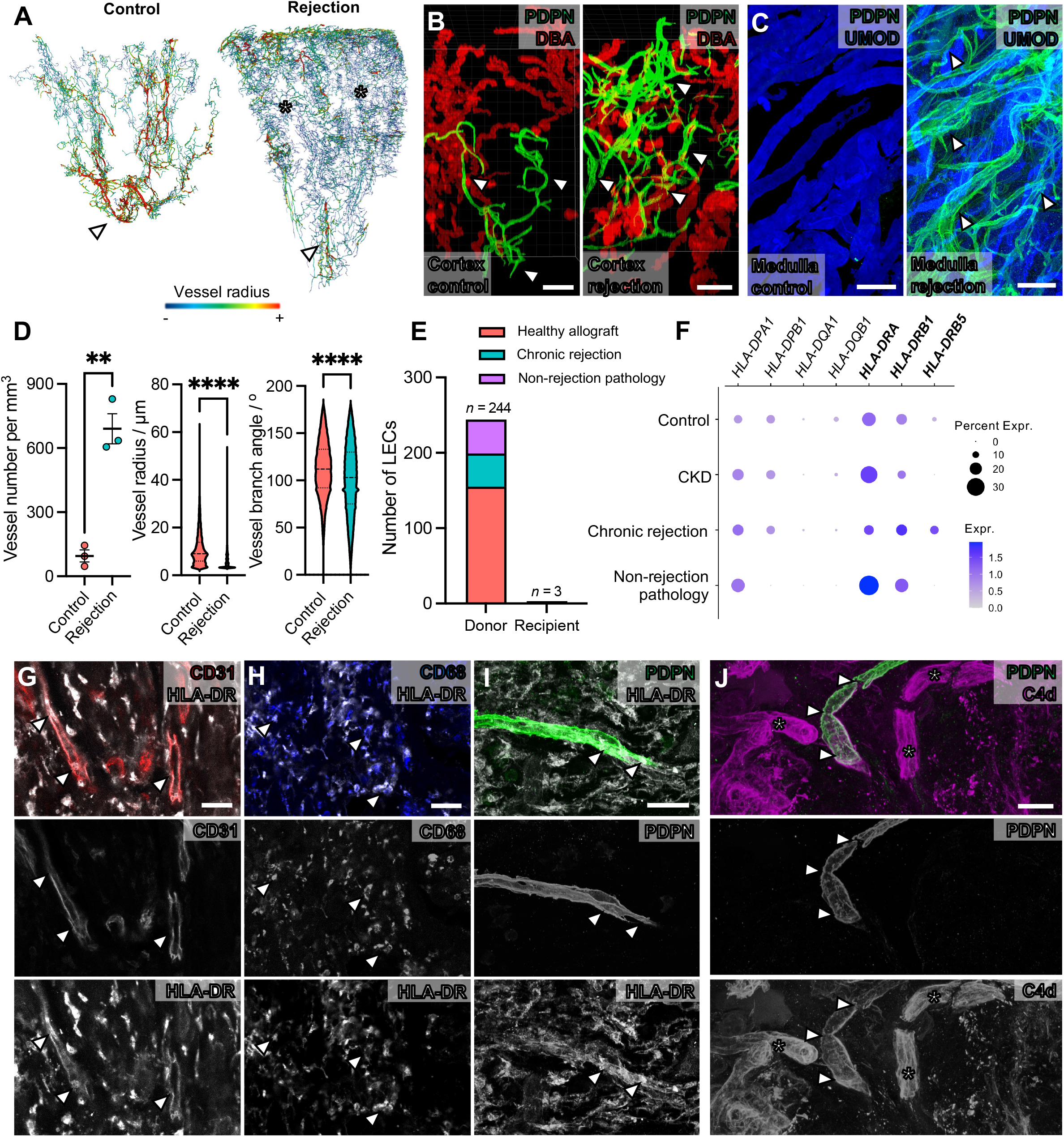
Mapping structure of lymphatics in chronic kidney transplant rejection and evidence for their targeting by alloantibodies. **(A)** 3D renderings LSFM imaging of kidney lymphatics in control (organ donor) and chronic transplant rejection tissues, representative of *n* = 3 images per group, where colour represents vessel branch radii, with blue representing the smallest radius (smallest radius in blue < 3.5 μm; asterisks, largest radius in red, > (>18 μm, arrowheads). **(B-C)** Confocal imaging of lymphatics (arrowheads) in the renal cortex, adjacent to DBA^+^ tubules (**B**) and medulla, adjacent to UMOD^+^ tubules (**C**), demonstrating the expansion of lymphatics in the cortex and their infiltration into the medulla. Images are representative of six regions of interest samples across *n* = 3 kidneys per condition. Scale bars = 200 μm (**B**), 100 μm (**C**). (**D**) Quantitative analysis of lymphatic vessel branch metrics in control and rejected kidneys. Data are presented either on the level of each kidney (scatterplot, *n =* 3 per group) or with all vessels from all kidneys pooled (violin plots, *n =* 75,036 vessel branches from donor *vs*. n = 1,048,576 vessel branches in rejection). Unpaired *t*-test revealed a significant increase in the number of vessels per unit volume in rejection, (95.12 ± 49.21 *vs*. 690.3 ± 121.6 vessels / mm^3^, *p* = 0.0014) whereas Mann-Whitney U tests revealed shifts in the distribution of vessel branch radius (*p* < 0.0001) and angle (*p* < 0.0001). (**E**) Donor or recipient status of the lymphatics in allograft tissues from the scRNA-seq atlas. Cells are grouped by healthy allograft, chronic rejection or non-rejection pathology of allografts (pyelonephritis, focal segmental glomerulosclerosis). (**F**) Dot plot of the expression of transcripts encoding MHC class II molecules within lymphatics, with enrichment of human leukocyte antigen (*HLA*)-*DRB5* expression in alloimmune rejection (log_2_FC = 2.05, *p* = 1.92 × 10^−7^). (**H-I**) 3D confocal images of HLA-DR expression (arrowheads) in PECAM1^+^ endothelia (**G**), CD68^+^ macrophages (**H**) and PDPN^+^ lymphatics (**I**). Images are representative of five regions imaged across *n =* 2 kidneys with chronic transplant rejection. (**J**) 3D Confocal images showing deposition of complement component C4d, representative of five regions imaged across *n =* 2 kidneys with chronic transplant rejection. C4d deposition is observed in PDPN^+^ lymphatics (arrowheads) and presumptive blood capillaries (asterisks). All scale bars = 30 μm.

This distortion of lymphatic architecture is reminiscent of the blood endothelial disruption observed in chronic rejection (Adair et al. 2007), where donor-derived endothelial cells expressing MHC Class II molecules human leukocyte antigen (HLA)-DR, HLA-DQ and HLA-DP (Kosmoliaptsis et al. 2014; Daniëls et al. 2020) are targeted by alloantibodies (Loupy and Lefaucheur 2018). To examine whether lymphatics might also be subject to alloantibody-mediated damage, we assessed the donor or recipient status of lymphatics derived from allografts with chronic rejection and compared this to healthy allografts and other non-rejection pathologies of the allograft including pyelonephritis and focal segmental glomerulosclerosis. Within our dataset, the majority of transplant lymphatic cells were donor-derived (*n =* 244/247, 98.8%), with a small contribution from recipient cells to lymphatics (*n =* 3/247, 1.2%), in line with a previous study of sex-mismatched renal allografts (Kerjaschki et al. 2006). To assess whether lymphatics might also be subject to alloantibody-mediated damage, we assessed lymphatic MHC Class II expression in the dataset. Kidney lymphatics expressed *HLA-DP* and *HLA-DR* transcripts, with little *HLA-DQ* detectable (**Fig.3F**), and *HLA-DRB5* expression was greater in kidneys with rejection than those with CKD (log_2_FC = 2.05, *p* = 1.92 × 10^−7^). As well as being evident in CD31^+^ blood endothelial cells (**Fig.3G**) and CD68^+^ macrophages (**Fig.3H**), the latter previously reported (Muczynski et al. 2003; Muczynski et al. 2001), HLA-DR protein expression was confirmed on PDPN^+^ lymphatics (**Fig.3I**). Consistent with alloantibody-mediated complement activation, we found C4d deposition on PDPN^+^ lymphatic vessels in the cortex of transplant rejection tissues (**Fig.3J**). Overall, our data suggest that allograft lymphatics may be directly targeted by donor-specific HLA antibodies leading to complement activation, in an analogous mechanism to that described for blood vascular endothelium, potentially leading to the distorted lymphatic architecture observed in rejecting allografts.

### Spatial and molecular analysis of lymphocyte trafficking by kidney lymphatics in chronically rejecting transplants

In homeostasis and inflammation, lymphatics secrete chemokines such as CCL21, generating gradients that promote leukocyte chemotaxis, entry into lymphatics and tissue efflux (Luther et al. 2002). Egress of leukocytes reduces local inflammation whilst simultaneously delivering these cells to draining lymph nodes, the sites where adaptive immune responses are generated (Johnson 2021; Steele and Lund 2021). Given the potential for lymphatics to influence alloreactive lymphocyte trafficking in chronic rejection, we investigated the localisation of PDPN^+^ lymphatics relative to CD20^+^ B cells and CD4^+^ T cells, central to the adaptive alloimmune response through recognition of differences between donor and recipient histocompatibility antigens by recipient T cells and production of antibodies by the B cell lineage (Duneton et al. 2022). We built on the findings of previous immunohistochemical studies describing the presence of lymphocytes within and around rejecting allograft lymphatics (Tsuchimoto et al. 2017; Kerjaschki et al. 2004) by quantifying intraluminal and extra-lymphatic tissue lymphocytes using 3D image analysis (**Fig.4A-B**). Luminal CD20^+^ B cell density was halved in rejecting allografts compared with controls (*p* = 0.0042) (**Fig.4C**), although the absolute number of B cells did not significantly differ (*p =* 0.977), suggesting a change in lymphatic volume rather than B cell abundance. Conversely, the total number of intra-luminal CD4^+^ T cells increased in chronic rejection (*p =* 0.0032), with the mean CD4^+^ T cell density increasing threefold (**Fig.4D**, *p <* 0.0001), higher than that of the surrounding allograft parenchyma (**Fig.4D**, mean *p <* 0.0001). To further examine the spatial interactions between lymphocyte subsets and lymphatics in transplant rejection, we assessed the normalised distance (Davis et al. 2017) of each lymphocyte to its nearest lymphatic vessel (**Fig.4E-F**). No significant spatial association was observed between CD20^+^ B cells and lymphatics in either control kidneys (*n =* 703 cells; *p =* 0.631) or rejecting allografts (*n =* 2,963 cells; *p =* 0.326) (**Fig.4G**). In contrast, CD4^+^ T cells (*n* = 2,149 cells across two controls) had a peak frequency within 0-100 μm from the nearest lymphatic vessel and showed greater spatial proximity to these vessels than would be expected if randomly distributed (*p =* 0.029). This spatial proximity of CD4^+^ T cells to lymphatics was lost in chronic transplant rejection (*n =* 4,382 cells, *p =* 0.699) (**Fig.4H**), suggesting dysregulated CD4^+^ T cell trafficking via lymphatics.

**Figure 4.**
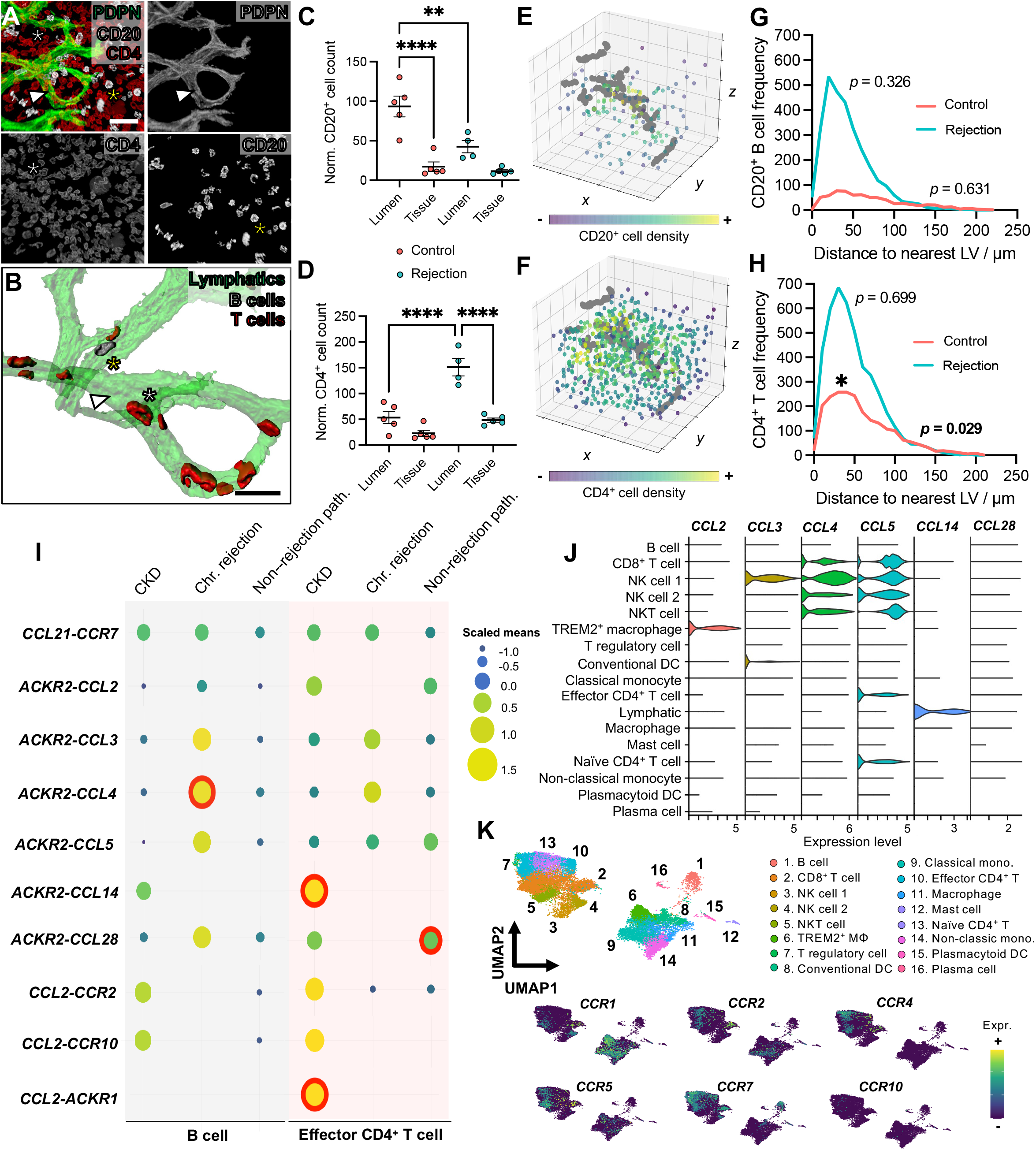
Inferring the spatial and molecular basis of lymphocyte trafficking by kidney lymphatics in alloimmunity. (**A & B**) Segmented (**A**) and rendered (**B**) confocal images of PDPN^+^ lymphatics (white arrow), CD20^+^ B cells (yellow asterisk) and CD4^+^ T cells (white asterisk). In **B**, the transparency of rendered lymphatics is increased to visualize intra-luminal B cells and T cells. Scale bars = 30 μm. (**C & D**) Number of intra-luminal CD20^+^ B cells (**C**) or CD4^+^ T cells (**D**), normalised by volume, was quantified and compared to that of the tissue parenchyma. Each point represents one volume of interest imaged. Luminal CD20^+^ B cell density in controls (organ donor) was significantly higher than that of the tissue parenchyma (mean difference = 76.2, 95% CI = 42.9-109.4, *p <* 0.0001), and approximately halved in rejecting allografts (mean difference = 50.8, 95% CI = 15.6-86.1, *p* = 0.0042). Intraluminal CD4^+^ T cell count was significantly higher in rejection (mean difference = 97.7, 95% CI = 52.8-142.7, *p <* 0.0001) and, within rejection, was higher than that of the surrounding parenchyma (mean difference = 102.7, 95% CI = 57.6-147.6, *p <* 0.0001). (**E & F**) Spatial point-patterns of peri-lymphatic CD20^+^ cell (**E**) or CD4^+^ cell (**F**) density, where lymphatic branch points represent grey dots and CD20^+^ cells are colour-coded according to their density around the lymphatic network. **(G & H)** Histograms of CD20^+^ cell (**G**) or CD4^+^ T cell (**H**) frequency as a function of distance from the nearest lymphatic vessel. *p* values demonstrate whether lymphocytes are clustered around lymphatics greater than would be expected under complete spatial randomness. The only significant association observed was between CD4^+^ T cells and lymphatics in control kidneys (*p =* 0.029). (**I**) CellPhoneDB dot plot of scRNA-seq data demonstrating top 10 chemokine interactions between lymphatics and B cells or effector CD4^+^ T cells, partitioned by disease aetiology. Dot size represents the scaled mean expression of the interaction, and those encircled with a red ring are deemed statistically significant by CellPhoneDB. **(J)** Violin plots of transcripts encoding atypical chemokine receptor 2 (ACKR2) ligands across all immune cell types within the chronic transplant rejection scRNA-seq dataset. **(K)** UMAP of all immune cell subsets with feature plots showing expression of receptors for ACKR2 ligands across these cell types. All imaging data is representative of *n* = 5 imaging volumes each acquired from *n =* 2 allografts with chronic rejection and *n =* 2 controls.

To investigate molecular signals underpinning lymphocyte chemotaxis by lymphatics, we analysed the single-cell transcriptomes of kidney lymphatics and immune cells in isolation, including B cells and effector CD4^+^ T cells, interrogating ligand-receptor interactions using CellPhoneDB (Efremova et al. 2020) (**Fig.S2A**). We detected ten predicted chemokine interactions between lymphatics and lymphocytes (**Fig.4I**), with the known lymphatic-leukocyte chemotactic cue mediated *via CCL21* signalling to its receptor CC receptor (*CCR)7* (Johnson 2021; Kerjaschki et al. 2004) evident across all groups. Lymphatics also expressed the atypical chemokine receptor 2 (*ACKR2*) that acts to sequester chemokines such as CCL2, CCL3, CCL4 and CCL5 and generate chemotactic gradients (Bonavita et al. 2016). These chemokines and their complementary receptors were varyingly expressed by macrophages and natural killer (NK), NKT, CD8^+^ T, CD4^+^ T and B cells in our scRNAseq data (**Fig.4J-K**). In rejection, there was a significant increase in the predicted interaction mediated by *ACKR2* on lymphatics and B cell-derived *CCL4* (**Fig.4I**). In contrast, a number of *ACKR2-*mediated interactions between lymphatics and effector CD4^+^ T cells were lost in rejection compared to control tissues and CKD, particularly *ACKR2-CCL2, -CCL14* and -*CCL28* (**Fig.4I**). These molecular analyses suggest a broad role for lymphatics in regulating kidney leukocyte dynamics. Attenuation of atypical chemokine interactions may underpin the disrupted localisation of CD4^+^ T cells observed in our 3D imaging of rejecting kidneys, analogous to the aberrant accumulation of immune cells within lymphatics present in ACKR2-deficient mice (K. M. Lee et al. 2011).

### Putative immunomodulation of *in situ* adaptive immune responses by lymphatics in alloimmunity

Whether lymphatics propagate or inhibit alloimmunity in transplantation is controversial (Wong 2020; Vass et al. 2009). On one hand, expansion of lymphatics and their proximity to immune cell aggregates has been associated with improved graft survival, potentially due to increased leukocyte clearance from the kidney (Tsuchimoto et al. 2017; Stuht et al. 2007; Pedersen et al. 2020), whereas other studies found increased lymphatics in kidneys with greater fibrosis and immune activation (Palin et al. 2013; Vass et al. 2012; Talsma et al. 2017; Lin et al. 2021; Thaunat et al. 2006). To investigate how adaptive immune responses within human kidney allografts might be modulated by lymphatics, we profiled tertiary lymphoid structures (TLS); aggregates of lymphocytes capable of *in situ* activation of antigen-specific T cells (Lee et al. 2022; Dorraji et al. 2020; Sato et al. 2020; Ichii et al. 2022; Meylan et al. 2022; Schröder et al. 1996; Lee et al. 2006) and observed in proximity to lymphatics in immunohistochemical studies of rejecting and non-rejecting allografts (Stuht et al. 2007; Kerjaschki et al. 2004; Tsuchimoto et al. 2017). In addition to observing PDPN^+^ lymphatics in close proximity to T and B cell-containing aggregates within the cortex of rejecting allografts (**Fig.5A**), we dissected the spatiotemporal relationship between TLS and lymphatics using CD21, a follicular dendritic cell marker, to identify TLS, and peripheral lymph node addressin (PNAd), to delineate high endothelial venules (HEV); only present in late-stage TLS (Ruddle 2016; Robson and Kitching 2020; Sato et al. 2021; Drayton et al. 2006; Alsughayyir et al. 2017; Motallebzadeh et al. 2012) (**Fig.5B**). We found PDPN^+^ lymphatics in TLS of all stages (*n =* 9/9, 100%), whereas HEVs were only present in half of TLS (*n =* 5/9, 55.6% *p =* 0.023). In these late-stage TLS, PDPN^+^ lymphatic vessel tips were localised more closely than HEVs to the CD21^+^ TLS core (**Fig.5C**, mean distance = 49.53 ± 23.83 μm *vs*. 109.6 ± 25.13 μm, 95% CI = 24.33-95.76, *p =* 0.0047). Furthermore, an interconnected network of PDPN^+^ lymphatics joined adjacent TLS, whereas PNAd^+^ HEVs were limited to the vicinity of TLS (**Fig.5D**). Lymphatics and HEV cells were also transcriptionally distinct (**Fig.5E**); lymphatics expressed a number of anti-inflammatory, pro-reparative transcripts (Paulsen et al. 2008; Belle et al. 2019; Liu et al. 2020), including *TFF3* and reelin (*RELN*). Conversely, HEV cells expressed molecules involved in leukocyte recruitment and activation such as *CXCL16* (Di Pilato et al. 2021), fractalkine (*CX3CL1*) (Garcia et al. 2013), CD40 (Elgueta et al. 2009) and interleukin 32 *(IL32*) (de Albuquerque et al. 2020) (**Fig.5F**). To further explore potential anti-inflammatory roles of lymphatics, we assessed immune stimulatory or inhibitory interactions between lymphatics and CD4^+^ T cell subsets, including effector, naïve and regulatory T cells. Co-inhibitory molecular interactions that dampen effector T cell responses (Anderson et al. 2016), including the ligands poliovirus receptor (*PVR*) and galectin 9 (*LGALS9*), dominated between lymphatics and CD4^+^ T cell subsets (**Fig. 5G**) or CD8^+^ T cells (**Fig.S2B**). The predicted immune-inhibitory interactions mediated by *PVR* and *LGALS9* were also present in CKD and non-rejecting transplant kidneys (**Fig.S2C-D**), but to a lesser extent than in chronic rejection (**Fig.5H**). We confirmed PVR expression by PDPN^+^ lymphatics in transplants with chronic rejection and found CD4^+^ T cells directly contacting PVR^+^ regions of lymphatic vessels (**Fig.5I**).

**Figure 5.**
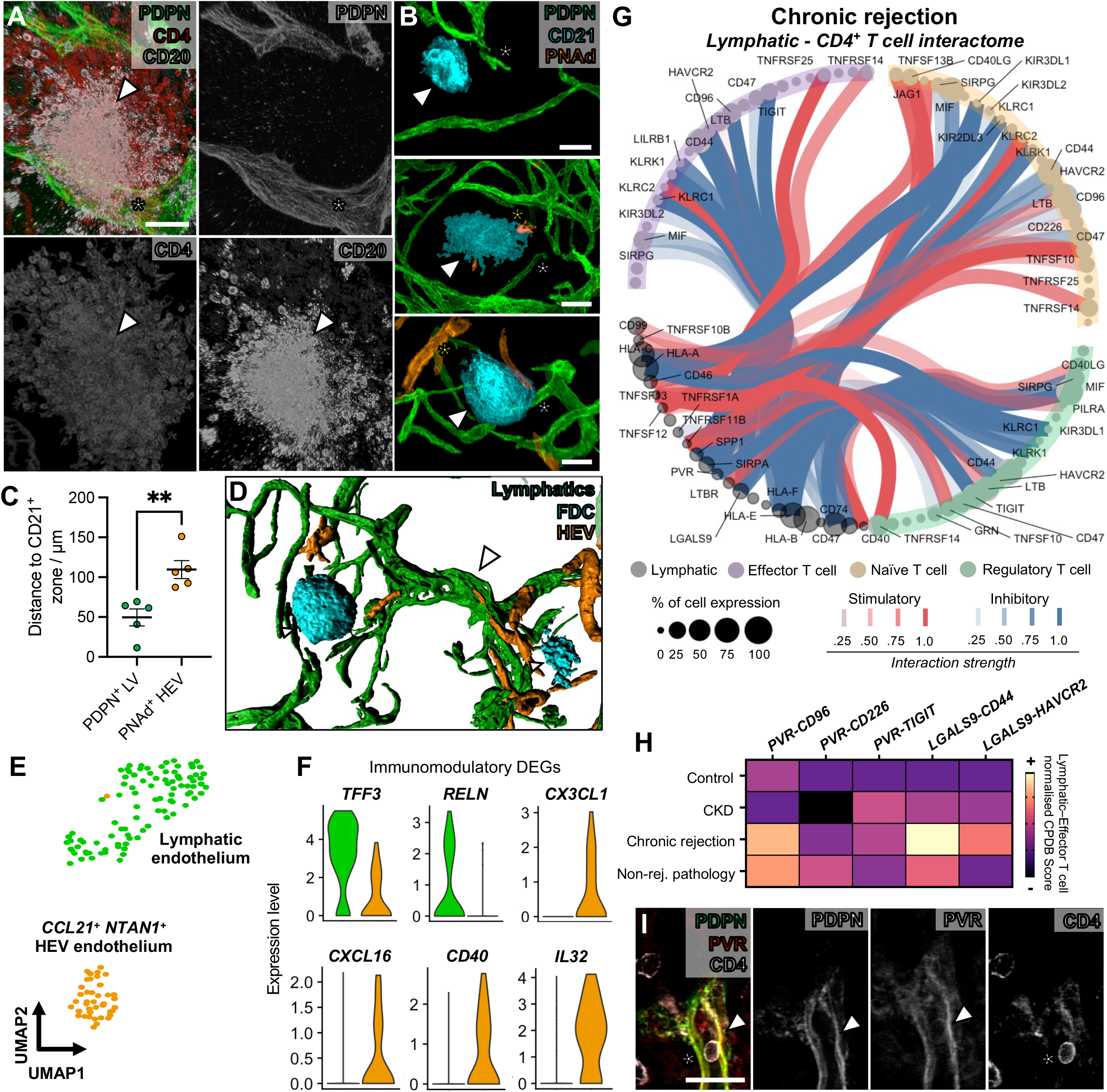
Interrogating the immunomodulatory molecular landscape of kidney lymphatics in alloimmunity. (**A**) Representative confocal images of PDPN^+^ lymphatics (white asterisk), CD20^+^ B cells and CD4^+^ T cells in regions with evidence of lymphocyte aggregation. A tertiary lymphoid structure (TLS) is shown (white arrowhead). Representative image of 4 T cell and B cell-rich TLS taken from *n =* 2 rejecting allografts. Scale bar = 40 μm. (**B**) Representative confocal images of TLS, containing PDPN^+^ lymphatics (white asterisk), CD21^+^ follicular dendritic cells (white arrowhead) and peripheral lymph node addressin (PNAd)^+^ high endothelial venules (HEV, yellow asterisk). 9 TLS were imaged across *n =* 3 rejecting allografts. Each image represents TLS at different stages, with either HEVs absent (early stage; top image), scant (mid-stage; middle image) or present (late-stage, bottom image). Scale bar = 50 μm. (**C**) 3D rendering of TLS interconnected by lymphatics, with the same molecular markers used in **B**. Such interconnections (white arrowhead) were observed between TLS in all (*n =* 3) rejecting allografts imaged. (**D**) Comparison of distance between the CD21^+^ TLS core and lymphatic vessel (green) or HEVs (orange), with each data point representing an individual TLS imaged. Lymphatic vessels were significantly closer to the CD21^+^ TLS core than HEVs (mean difference = 60.04, 95% CI = 24.33-95.76, *p =* 0.0047). **(E)** UMAP of lymphatics and blood vasculature expressing *CCL21* and *PNAd* within rejecting allografts in the scRNA-seq dataset. **(F)** Violin plots showing immunomodulatory candidates differentially expressed between lymphatic endothelium and HEV cells across rejecting allograft scRNA-seq data. Pro-reparative candidates (*TFF3, RELN*) are enriched in lymphatic endothelium whereas immune activating candidates (*CX3CL1, CXCL16, CD40, IL32*) are enriched in *CCL21*^*+*^ *PNAd*^*+*^ blood endothelium. **(G)** Circle plot ‘interactome’ of scRNA-seq data from rejecting allografts, including curated CellPhoneDB interactions derived from lymphatics and acting on different CD4^+^ T cell subsets. Each node represents a putative ligand or receptor, with node colour as cell type. Each line represents an interaction (stimulatory interactions = red, inhibitory interactions = blue. The size of the node represents the proportion of cells expressing the ligand or receptor, and the darkness of the line represents the strength of the interaction. **(H)** Heatmaps of immune checkpoint interactions detected between lymphatics and effector CD4^+^ T cells within the scRNAseq data across different aetiologies of disease, with colour representing the normalised CellPhoneDB score. **(I)** Imaging validation of expression of poliovirus receptor (PVR) in PDPN^+^ lymphatics (white arrowhead) and a contacting CD4^+^ T cell (white asterisk) from *n =* 2 rejecting allografts. Scale bar = 30 μm.

Collectively, 3D imaging and scRNA-seq analyses implicate lymphatics as an early and ubiquitous feature of lymphoid aggregates in kidneys with chronic rejection, forming an interconnecting vascular network between TLS. Lymphatics display a distinct molecular profile from their blood vascular counterparts in chronic rejection, expressing immune inhibitory molecules with the potential to regulate local alloreactive CD4^+^ T cells. These findings, akin to those recently delineated for mouse tumour (Gkountidi et al. 2021; Steele et al. 2022), central nervous system (Hsu et al. 2022) and dermal lymphatics (Churchill et al. 2022), suggest lymphatics modulate adaptive immune responses within the tissue microenvironment, beyond their roles in leukocyte egress; implicating these vessels as an emerging therapeutic target in chronic transplant rejection (Gupta et al. 2012; Sun et al. 2021).

## CONCLUSION

Overall, we have used 3D imaging and scRNA-seq to comprehensively characterise the human kidney lymphatic vasculature, providing spatial and molecular references for studies of kidney physiology and disease. We also provide new insights into the potential roles of lymphatics in transplant biology and immunity, pointing towards putative alloantibody targeting of lymphatics, accompanied by loss of lymphatic hierarchy and impairment of CD4^+^ T cell lymphatic trafficking as cellular hallmarks of chronic transplant rejection. These findings potentially explain why loss of graft function occurs despite lymphangiogenesis. Finally, we decipher relationships between kidney lymphatics and alloimmune responses, identifying that these vessels colonise and interconnect tertiary lymphoid niches and may partake in immunoregulatory crosstalk involving co-inhibitory checkpoint molecules. Such findings provide impetus to consider lymphatics as a key player in alloimmunity with therapies modulating lymphatics having potential to promote transplant longevity.

## ACKNOWLEDGEMENTS

The authors would like to acknowledge Professors Lucy Walker, Mark Lythgoe, Alan Salama (UCL) and Dr René Hägerling (Charité Universitätsmedizin Berlin) for their ongoing support and valuable discussions about the work, and Dr Kelvin Tuong (University of Cambridge) for producing scripts for visualisation and analysis of ligand-receptor pair analysis of scRNA-seq data (https://github.com/zktuong/ktplots). Surgical explantation was performed by the Renal Transplant surgical team at Royal Free Hospital, London, UK. Confocal microscopy was performed at the Light Microscopy Core Facility at UCL GOSICH and LSFM was performed at the UKRI Dementia Research Institute. All work was performed with the support of a grant from Kidney Research UK (IN_012_20190306), a Rosetrees Trust PhD Plus Award (PhD2020\100012) and Foulkes Foundation Fellowship to Dr Daniyal Jafree, a Wellcome Trust Investigator Award (220895/Z/20/Z) to Professor David Long, the National Institute for Health Research (NIHR) Biomedical Research Centre at Great Ormond Street Hospital for Children NHS Foundation Trust and University College London. All authors declare no conflicts of interest.

## AUTHOR CONTRIBUTIONS

DJJ, MRC, RM and DAL were involved in the conception of the study. Acquisition of material and laboratory experiments were performed by DJJ, MKJ, LGR, LMR, WJM, BIL, LW, TA, SM, JC, KC and KP. Analysis of 3D images or scRNA-seq data was performed by DJJ, BS, BD, HM, DM, NH, CW and SWS. Histopathological analysis and acquisition of clinical data was performed by LH and RM. Project oversight and supervision was provided by PJDW, MAB, PJS, MRC, RM, ASW and DAL. DJJ wrote the first draft of the paper, refined by DAL and MRC, and subsequently all authors were involved in revision and preparation of the final manuscript for submission.

## FIGURES AND FIGURE LEGENDS

**Supplementary Figure 1.**
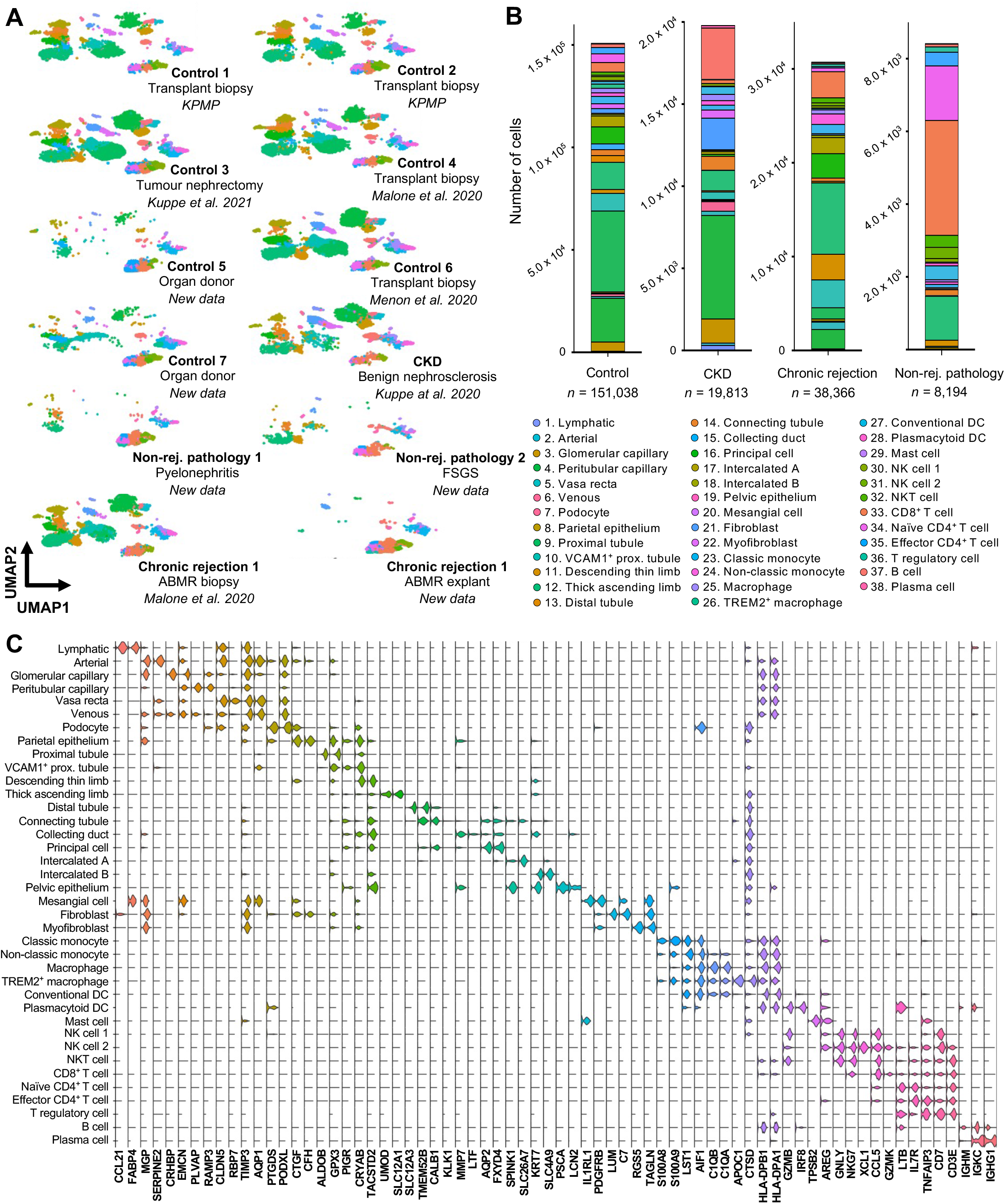
Generating a human kidney single-cell transcriptomic atlas featuring health and pathologies. (**A**) Uniform manifold approximation and projection (UMAP) of the kidney cell atlas coloured by cell type and partitioned by dataset. Data sources are included under each sample label with either appropriate references, new samples or whether data was derived from the Kidney Precision Medicine Project (KPMP). Samples included non-rejection ‘healthy’ transplant biopsies (Control 1: 7,259 cells, Control 2: 9,785 cells, Control 4: 38,850 cells and Control 6: 22,592 cells), non-tumorous regions of tumour nephrectomies (Control 3: 58,934 cells, chronic kidney disease due to benign nephrosclerosis (CKD): 58,934 cells), declined organ donor tissues (Control 5: 3,877 cells, Control 7: 9,741 cells) and surgically explanted allografts (Non-rejection pathology 1 (chronic pyelonephritis): 6,063 cells, Non-rejection pathology 2 (focal segmental glomerulosclerosis): 2,131 cells, Chronic rejection 1: 34,067 cells, Chronic rejection 2: 4,299 cells). (**B**) Stacked bar charts representing the relative proportions of each annotated cell type across the four study groups). (**C**) Stacked violin plot showing top two differentially expressed marker genes (*y* axis) by cell type (*x* axis), calculated using Seurat FindAllMarkers function.

**Supplementary Figure 2.**
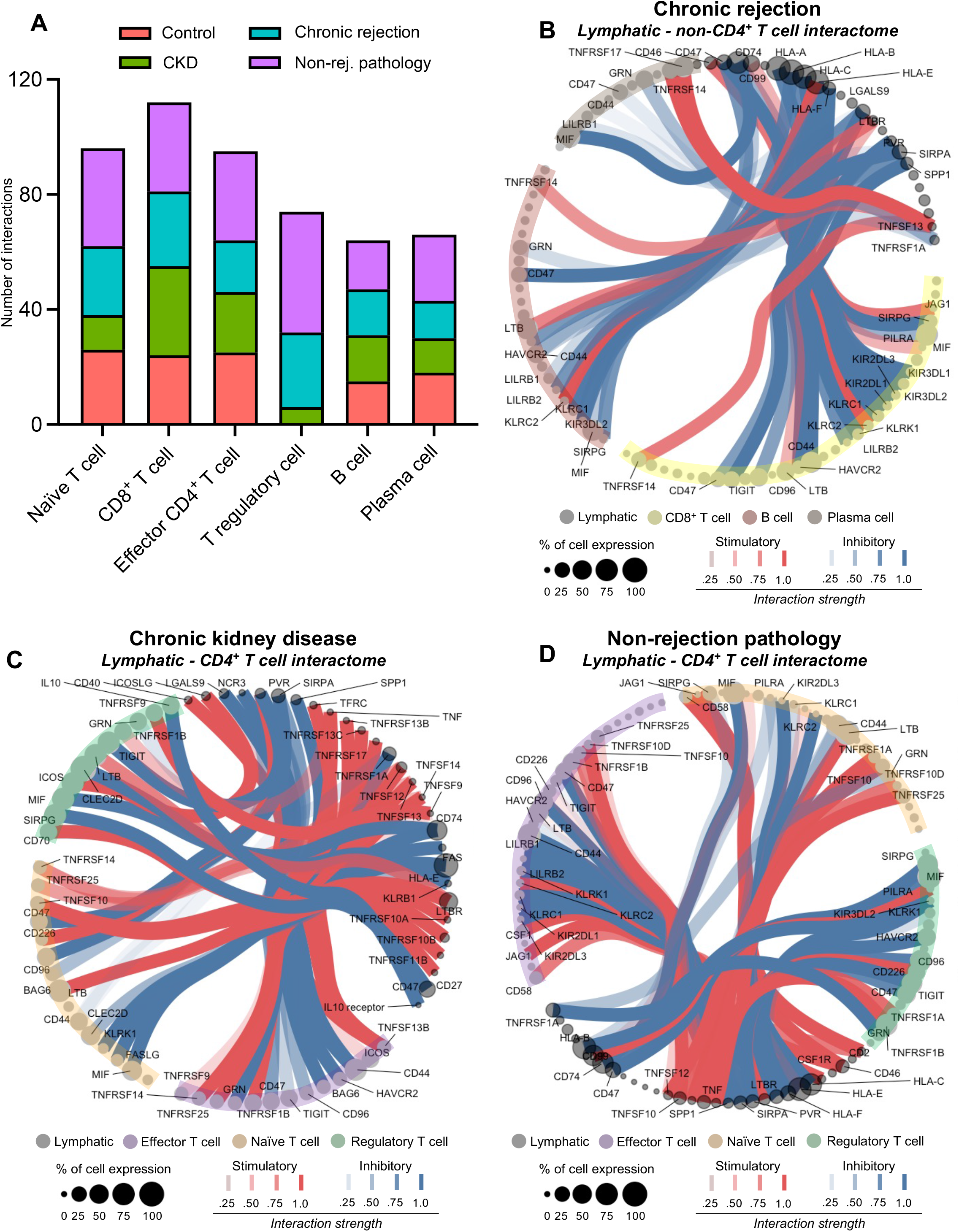
Cell-cell interactome landscape of lymphatics and adaptive immune subsets across kidney pathologies. (**A**) Bar chart showing the total number of CellPhoneDB-computed cell-cell interactions between lymphatics and adaptive immune cell subsets within single-cell RNA sequencing (scRNA-seq) data of control, chronic kidney disease (CKD), alloimmune rejection and non-alloimmune rejection tissues. (**B-D**) Circle plot of the transcriptional ‘interactome’, or putative cell-cell communication, between lymphatics and distinct immune cell subsets identified in the scRNA-seq dataset, computed between lymphatics and non-CD4^+^ T cells in alloimmune rejection (**B**), lymphatics and CD4^+^ cell subsets in CKD (**C**), and lymphatics and CD4^+^ cell subsets in non-rejection pathology (**D**). Each node represents a putative ligand or receptor, with node colour as cell type. Each line represents an interaction, with stimulatory interactions coloured in red and inhibitory interactions coloured in blue. The size of the node represents the proportion of cells expressing the ligand or receptor, and the darkness of the line represents the strength of the CellPhoneDB interaction.

## MATERIALS AND METHODS

### Three-dimensional imaging and analysis of human kidney lymphatics

#### Acquisition, fixation and storage of human tissue for three-dimensional imaging

Human adult kidney tissue was derived from four deceased patients who had opted in for organ donation and undergone multi-organ procurement, but for whom the kidneys had ultimately been declined for implantation by recipient transplant centres. Kidneys were retrieved by a UK National Organ Retrieval Services teams. Following *in situ* flushing of the abdominal organs with University of Wisconsin (UW) solution, the kidneys were removed and stored in UW at 4°C. Consent for the use of the organs for research was obtained from the donor family by Specialist Nurses in Organ Donation before organ retrieval and were then offered for research by NHS Blood & Transplant (NHSBT) if they were found to be unsuitable for transplantation by the surgical team. Ethical approval was granted by the National Research Ethics Committee in the UK (21/WA/0388) and was approved by The Royal Free London NHS Foundation Trust-UCL Biobank Ethical Review Committee (RFL B-ERC; NC.2018.010; IRAS 208955). Kidney allograft samples were obtained from three patients at Royal Free London NHS Trust undergoing nephrectomy for graft intolerance syndrome (*n* = 2) and graft malignancy (*n* = 1). Ethical approval was covered by a prior agreement (NC.2018.007, UCL Biobank Ethical Review Committee, Royal Free London NHS Foundation Trust, B-ERC-RF). All explants were performed by the transplant surgical team. Prior to acquisition, all patients were confirmed negative for COVID-19 by means of a qPCR test. After explant, pseudo-anonymised human adult kidney tissues were incubated overnight in Belzer University of Washington Cold Storage Solution (Bridge to Life Europe, London, UK) at 4°C. Prior to fixation, human adult kidney was manually dissected into ∼3mm full-thickness sub-regions containing cortex and outer medulla. These tissues were then incubated in 4% (w/v) paraformaldehyde (PFA, Sigma Aldrich), made up in 1 X phosphate buffered saline (PBS), at 4° C overnight. After fixation, all biological tissues were washed and stored in 1 X PBS with 0.02% (w/v) sodium azide to prevent contamination. Randomly selected pieces of human adult kidney were transferred to and stored in 70% ethanol for histology

#### Wholemount immunofluorescence

A modified version of the SHANEL protocol (Zhao et al. 2020) was implemented for wholemount immunolabelling of kidney tissues. Unless otherwise stated, steps were performed at room temperature, and reagents purchased from Sigma Aldrich. Tissues were dehydrated in a methanol series (50, 70%) in double distilled (dd)H2O, for one hour per step, before bleaching in absolute methanol with 5% (v/v) of 30% hydrogen peroxide solution overnight at 4°C. Thereafter, tissues were rehydrated in the methanol series, followed by incubation in 1 x PBS for one hour. Overnight incubation of tissues was performed in a 0.5 M solution of acetic acid at 4°C, followed by five hours of incubation at 4°C with 4 M guanidine hydrochloride, 0.05 M sodium acetate and 2% (v/v) Triton X-100 made up in PBS. Tissues were then permeabilised with 5% (w/v) solution of 3-((3-cholamidopropyl) dimethylammonio)-1-propanesulfonate (CHAPS) made up in ddH2O overnight. Then tissues were incubated for one day in blocking solution, comprising 1 x PBS with 0.2% Triton X-100, 5% (v/v) donkey or goat serum, 5% (v/v) pooled human plasma (Biowest, Nuaillé, France) and 10% (v/v) dimethyl sulfoxide (DMSO) before incubation in antibody solution (1 x PBS with 0.2% (v/v) Tween-20, 0.1% (v/v) of a 10mg/ml heparin solution in ddH2O, 0.1% (w/v) saponin, 2.5% donkey or goat serum, 2.5% pooled human plasma with primary antibodies at the appropriate concentration at 4°C. Blocking and antibody solutions were further supplemented with 1:150 Human TruStain FcX™ Fc Receptor Blocking Solution (BioLegend, London, UK), to reduce non-specific binding. Primary antibodies were incubated for 3-4 days, before replenishing the antibody solution and re-incubation for 3-4 days. Subsequently, tissues were washed in 1 x PBS with 0.2% Tween-20 four times for 1 hour per wash, before incubation in antibody solution with secondary antibodies at 1:200 at 4°C for four days. Tissues were then washed again in 1 x PBS with 0.2% Tween-20 four times for 1 hour each and stored until dehydration and clearing.

#### Primary antibodies, lectins and secondary antibodies

In order of appearance in the manuscript, the following primary antibodies or lectins were used in 1.5ml incubations at the indicated concentrations: mouse anti-PDPN monoclonal (clone: D2-40, 1:100, M3619, Aligent), rabbit anti-PROX1 polyclonal (1:200, ABN278, Merck), goat anti-LYVE1 polyclonal (1:100, AF2089, R&D Systems), fluorescein-conjugated LTL (1:50, Vector Laboratories), rabbit anti-UMOD monoclonal (clone: EPR20071, 1:100, ab207170, Abcam), fluorescein-conjugated UAE-I (1:50, Vector Laboratories), rabbit anti-LRP2 polyclonal (1:50, ab76969, Abcam), rabbit anti-CALB1 monoclonal (clone: EP3478, 1:100, ab108404, Abcam), rhodamine-conjugated DBA (1:50, Vector Laboratories), mouse anti-CDH1 monoclonal (clone: HECD-1, 1:50, ab1416, Abcam), mouse anti-PECAM1 monoclonal (clone: JC70A, 1:50, M0823, Dako), mouse anti-CD68 monoclonal (clone: KP1, 1:100, ab955, Abcam), rabbit anti-αSMA polyclonal (1:50, ab5694, Abcam), rabbit anti-HLA-DR monoclonal (clone: EPR3692, 1:100, ab92511, Abcam), rabbit anti-C4d polyclonal (1:100, 0300-0230, Bio-Rad), rabbit anti-CD4 monoclonal (clone: EPR6855, 1:100, ab133616, Abcam), goat anti-CD20 polyclonal (1:100, ab194970, Abcam), goat anti-PVR polyclonal (1:100, AF2530, R&D Systems), rabbit anti-CD21 monoclonal (clone: EPR3093, 1:200, ab75985, Abcam), rat anti-PNAd monoclonal (clone: MECA-79, 1:100, MABF2050, Sigma). All secondary antibodies were purchased from ThermoFisher Scientific, were conjugated to AlexaFluor fluorophores (488, 546, 568, 633 or 647) and were used at a concentration of 1:200 of the original secondary antibody stock. Controls for each panel involved omission of the primary antibody and including the secondary antibody only.

#### Solvent-based optical clearing

Tissues were dehydrated in a methanol series (50%, 70%, 100%) for 1 hour per step. BABB (benzyl alcohol and benzyl benzoate in a 1:2 ratio), was used for clearing, with all solutions containing BABB kept in glass scintillation vials (VWR International, Lutterworth, UK). Clearing was performed in glass scintillation vials, first using BABB:methanol in a 1:1 ratio, and thereafter BABB alone, until samples equilibrated and achieved transparency.

#### Confocal microscopy

We took advantage of the z-depth achievable by upright confocal microscopy whilst protecting the microscope objectives. All tissues were placed between a large coverslip and cover glass, supported by a O-Ring (Polymax Ltd, Bordon, UK) made from BABB-resistant rubber, as described previously (Jafree et al. 2020). Confocal images were acquired on an LSM880 upright confocal microscope (Carl Zeiss Ltd.), with a 2.5x/numerical aperture (NA) 0.085 Pan-Neofluar Dry objective (working distance; WD = 8,800 μm) for low-resolution imaging, and 10×/NA 0.5 W-Plan Apochromat water dipping objective (working distance; WD = 3,700 μm) for high-resolution imaging. Gallium arsenide phosphide (GaAsP) internal and external detectors were used for high sensitivity. To obtain higher resolution imaging, an Airyscan setting (Huff 2015), consisting of a 32-channel (GaAsP) photomultiplier tube area detector.

#### Lightsheet fluorescence microscopy

3D imaging of cleared tissues was performed using a custom-built mesoscale selective plane illumination microscope (mesoSPIM) (Voigt et al. 2019). The cleared tissue was secured in a 3D-printed holder and immersed in BABB solutions inside a quartz cuvette (40 × 40 × 100 mm). Fluorescence images were acquired with an Olympus MVX-10 macroscope at 1x magnification, resulting in a voxel size of 6.55 × 6.55 × 5 μm^3^. PDPN fluorescence signals were obtained using 638 nm laser excitation and 633nm long-pass optical filtering of emitted light, while autofluorescence was captured using 488 nm laser excitation and a 520/35 nm bandpass emission filter. Lightsheet illumination from both sides of the cuvette was carefully aligned after the sample was positioned at the centre of the macroscope’s field of view and delivered simultaneously to capture a single *z*-stack image.

#### Post-acquisition image processing

All images were then exported to FIJI (NIH, Bethesda, US). Confocal image stacks were separated into individual fluorescence channels, and the Despeckle and Sharpen tools were used to reduce non-specific background fluorescence. Where maximum intensity *z-* projections or optical *z-*sections were required, scale bars were applied and images and exported as TIFF files.

#### Image visualisation and binarization of three-dimensional imaging data

Visualisation of confocal 3D reconstructions were performed by importing confocal images to the commercial software, Imaris (v8.2, Bitplane). The Isosurface Rendering tool in Imaris allows the extraction of surfaces based on fluorescence intensity. This was used to generate segmented images fluorescence masks to better visualise expression patterns, or to generate binarized outputs for extraction of quantitative vessel branching metrics. LSFM data was imported into Amira (v2020.2, Fisher Scientific) and the vasculature segmented using intensity thresholding and region growing using the Magic Wand tool to generate a binarized network.

#### Extraction of vessel branching metrics from three-dimensional imaging data

Segmented and binarized confocal and LSFM images were imported as TIFF image stacks into Amira. The Filament Editor tool was used in Amira to generate spatial statistical parameters including vessel branch number, lengths, diameter and volumes from each segmented lymphatic plexus. The resulting values were exported these as CSV files.

#### Spatial statistical analysis of lymphatic-lymphocyte relationships

Lymphatic 3D-skeletons were extracted from binarised confocal stacks using the BoneJ Skeletonise3d function in FIJI (Lee et al. 1994). CD4^+^ T cell and CD20^+^ B cell counts, centroids and areas were obtained using 3d-objectcounter with no further pre-processing (Bolte and Cordelières 2006). The mean distance of each cell from the nearest point of the lymphatic network (*d*) was calculated using the cross-product 3D point-line distance:

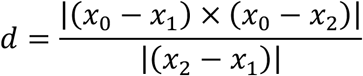

where *x*_1_ and *x*_2_ are the two closest adjacent nodes from the lymphatic 3D skeleton; found by minimizing cross-nearest neighbor distances, and *x*_0_ is the centroid of the cell of interest. To evaluate whether the cell distances were different from what would be expected by chance, within each region of interest, the CD4^+^ T cell and CD20^+^ B cell populations were randomly redistributed under complete spatial randomness for twenty simulations. A comparison was then made as to whether the measured mean cell-lymphatic distances fell within the 95% confidence intervals obtained through the simulations under complete spatial randomness.

### Single-cell transcriptomic analysis of human kidney lymphatics

#### Acquisition of material for single-cell transcriptomics and generation of a human kidney cell atlas

The scRNA-seq dataset from this study consisted of previously published data and five new samples. New samples were collected under the NHSBT service (REC12/EE.0446) and included two control biopsies from kidneys donated for transplantation but deemed unsuitable for use due to renal vein thrombus or mycotic pseudoaneurysm, and three further pathological samples including one chronic rejection sample (antibody-mediated) and two samples with non-rejection pathologies (chronic pyelonephritis or focal segmental glomerulosclerosis). These tissues were processed for scRNA-seq using the 10X Genomics Chromium system as previously described (Stewart et al. 2019). Briefly, samples were manually dissected into small pieces and digested for 30 minutes at 37°C, in 25 μg/ml Liberase TM (Roche) and 50 μg/ml DNase (Sigma) in 5 ml RPMI (Gibco) using gentleMACS (Miltenyi Biotec). The suspension was passed through a 100 μm cell strainer (Falcon), washed with PBS, and enriched for live cells using a Dead Cell Removal kit (Miltenyi Biotec) before washing with PBS. Cells were loaded according to the protocol of the Chromium single cell 3’ kit (v3 Chemistry). Sequencing was performed on an Illumina Hiseq 4000. Raw reads from samples and previously published scRNA-seq data of the human kidney, including samples from non-tumorous regions of tumour nephrectomies with or without CKD (Kuppe et al. 2021), live allograft biopsies with or without antibody-mediated rejection (Malone et al. 2020), transplant biopsies from the Kidney Precision Medicine project (https://atlas.kpmp.org/repository/) were mapped and quantified using cellranger software (10X Genomics). Data then underwent quality control, normalisation, feature selection and dimension reduction using the Scanpy package in Python (Wolf et al. 2018). Integration of samples from different batches was performed using scvi-tools (Lopez et al. 2018) before semi-automated cell type annotation of clusters using CellTypist (https://www.celltypist.org). The data was then converted using seurat-disk before further analysis using the Seurat package (Hao et al. 2021) in R. Unless otherwise stated, all downstream steps were performed in Seurat.

### Differential expression analysis

The *FindAllMarkers* function was used for differential expression analysis. Wilcoxon Rank Sum tests were used to assess statistically significant (adjusted *p* value ≤ 0.05) between average log fold change values of expression. Selected differentially expressed genes were visually represented using the *VlnPlot* function or *DoHeatMap* functions.

### Gene ontology analysis

Gene ontology (GO) analysis was performed using the PANTHER tool for gene classification (Mi et al. 2021). Lists of differentially expressed genes, were exported and input into the PANTHER web tool (v16.0, http://www.pantherdb.org), using statistical overrepresentation tests to group genes using the GO biological processes complete database. Fisher’s Exact tests were used to assess for statistical enrichment of genes for selected GO terms, and a false discovery rate (FDR) p ≤ 0.05 was considered significant.

### CellPhoneDB

To infer putative cell-cell interactions in single-cell RNA sequencing data, the CellPhoneDB resource (Efremova et al. 2020) was used. Using normalised count and metadata files obtained from Seurat, CellPhoneDB was called by running appropriate commands, obtained from https://github.com/Teichlab/cellphonedb, in the command line through a Python virtual environment. The *statistical_analysis* method was used to assess predicted interactions, before functions in ktPlots (https://github.com/zktuong/ktplots) were used to generate custom dot plots or circle plots.

### Statistical analysis and data presentation

#### Sample size estimation

In prior work examining the 3D architecture of lymphatic vessels in lymphangiomatous skin biopsies, conclusions were drawn based on the evaluation of three samples within the control group (Hägerling et al. 2017), and so a minimum of three patients per group were used to draw conclusions. For scRNA-seq, the number of samples and cells to be analysed was limited by the size of the dataset. The specific number of replicates used for each experiment and the number of regions images are indicated in the figure legends,

#### Reproducibility

Descriptive conclusions are drawn based on a minimum of four imaging volumes of interest, each taken from samples from at least two different human kidneys. The annotated and processed Seurat object, along with the scripts used for imaging analysis and interrogation of scRNA-seq data will be made publicly available upon publication of the manuscript.

#### Data presentation

All confocal and brightfield images were exported and saved as TIFF files. Where brightness or contrast were adjusted, this was applied uniformly across all conditions within the same figure, and details are stated in figure legends. Graphs were generated in GraphPad PRISM and saved as TIFF format. Visualisations from scRNA-seq analysis were performed in RStudio and PNG screenshots were taken and saved. Figures were compiled in Microsoft PowerPoint (Microsoft, Redmond, US) and saved as PDF format.

#### Statistics

Except for scRNA-seq analysis and lymphatic-lymphocyte spatial relationships, all remaining statistical comparisons were performed using GraphPad PRISM. A two-tailed *p* value of less than 0.05 was considered statistically significant. For continuous data, Shapiro-Wilk tests were used to assess normality of distribution and Brown-Forsythe tests were used equality of variance. Where normal distribution and equality of variances were satisfied, data is presented as mean ± standard deviation. When graphed, error bars were used to represent the standard error of the mean. Student’s *t*-test was used to compare two groups and ANOVA was used to compare more than two groups, applying post-hoc Bonferroni tests to provide adjusted *p* values for multiple comparisons. Statistics for scRNA-seq analysis were performed in RStudio and are as detailed above.

