## Supplementary Table 1 for "Three-dimensional imaging and single-cell transcriptomics of the human kidney implicate perturbation of lymphatics in alloimmunity"

| ID | DONOR DEMOGRAPHICS | | | | RENAL FUNCTION | | ORGAN CHARACTERISTICS | | |
|---|---|---|---|---|---|---|---|---|---|
| | Age | Sex | Ethnicity | Cause of death | Serum creatinine | eGFR | Type of organ | Reason for decline | Histological report |
| **NK1** | 68 | M | Cauc. | Intraparenchymal haemorrhage | 74 | 90 | DBD | Cancerous lesions on contralateral kidney | Chronic damage <2%; 7/115 obsolete glomeruli; mild tubular blebbing; mild fibro-intimal proliferation of vessels |
| **NK2** | 61 | M | Cauc. | Acute cerebral infarction | 94 | 75 | DCD | Severe aortic atherosclerosis | Chronic damage <10%; 5/60 obsolete glomeruli; occasional tubular flattening, blebbing and vacuolation; mild fibro-intimal proliferation of vessels |
| **NK3** | 57 | M | Cauc. | Hypoxic brain injury | 60 | 90 | DCD | Severe aortic atherosclerosis | Chronic damage <10%; mild lymphocytic infiltrate; 7/80 obsolete glomeruli. occasional occasional tubular flattening, blebbing and vacuolation; mild focal fibro-intimal proliferation of vessels |
| **NK4** | 22 | M | Cauc. | Traumatic head injury | 89 | 90 | DCD | Hydronephrosis and PUJ obstruction | Chronic damage ~0%; 0/69 obsolete glomeruli; mild tubular flattening; mild fibro-intimal proliferation of vessels |

**Table S1. Clinicopathological information for organ donors from which control tissues were derived for 3D imaging.** Age in years, serum creatinine in μmol/L, estimated glomerular filtration rate (eGFR) in ml/min/1.73m$^2$. Cauc., Caucasian; DBD, donor after brainstem death; DCD, donor after cardiac death; PUJ, pelvic ureteric junction

| ID | DONOR DEMOGRAPHICS | | | TRANSPLANT CHARACTERISTICS | | | | REJECTION INFORMATION | | |
|---|---|---|---|---|---|---|---|---|---|---|
| | Age | Sex | Ethnicity | Year of transplant | Type of organ | HLA mismatches | Drugs received | Indications for explant | Donor-specific antibodies | Histological report |
| **CR 1** | 41 | M | Cauc. | 2006 | Cadaveric | HLA-A (2)<br>HLA-B (1)<br>HLA-DR (1) | MMF tacrolimus | Failing transplant and two cancerous lesions | Anti-HLA-DQ5 (10778 MFI) | IF/TA present; glomerulosclerosis; moderate fibro-intimal proliferation of vessels with lymphocyte invasion; hyaline arteriosclerosis; tubulitis; interstitial lymphoplasmacytic infiltrate. Features of both TCMR and ABMR. |
| **CR 2** | 79 | F | Asian | 2007 | Live | Unknown | MMF cyclosporine | Failing transplant and graft intolerance | Unknown | IF/TA present; ischaemic glomeruli; peritubular capillaritis, endarteritis and chronic vasculopathy; tubilitis; interstitial lymphoplasmacytic infiltrate. Features of both TCMR and ABMR. |
| **CR 3** | 76 | F | Asian | 1995 | Cadaveric | HLA-A (1)<br>HLA-B (1)<br>HLA-DR (0) | Cyclosporine | Failing transplant and pain over graft | Anti-HLA-A2 (MFI 19807) Anti-HLA-A31 (MFI 25896) Anti-HLA-B60 (MFI 24681) | IF/TA present; mostly obsolete glomeruli; severe fibro-intimal proliferation of vessels with endarteritis; interstitial lymphoplasmacytic infiltrate; calcification and simple cysts present. Features of both TCMR and ABMR present. |

**Table S2. Clinicopathological information for patients from whom transplant rejection tissues were derived for 3D imaging.** Age in years. Cauc., Caucasian; DBD, donor after brainstem death; DCD, donor after cardiac death; HLA, human leukocyte antigen; MFI, mean fluorescence intensity; MMF, mycophenolate mofetil.
